# Analysis of HIV-1 latent reservoir and rebound viruses in a clinical trial of anti-HIV-1 antibody 3BNC117

**DOI:** 10.1101/324509

**Authors:** Yehuda Z. Cohen, Julio C. C. Lorenzi, Lisa Krassnig, John P. Barton, Leah Burke, Joy Pai, Ching-Lan Lu, Pilar Mendoza, Thiago Y. Oliveira, Christopher Sleckman, Katrina Millard, Allison L. Butler, Juan P. Dizon, Shiraz A. Belblidia, Maggi Witmer-Pack, Irina Shimeliovich, Roy M. Gulick, Michael S. Seaman, Mila Jankovic, Marina Caskey, Michel C. Nussenzweig

**Affiliations:** Laboratory of Molecular Immunology, The Rockefeller University, New York, NY 10065, USA; Department of Physics and Astronomy, University of California, Riverside, CA 92521, USA; Division of Infectious Diseases, Weill Cornell Medicine, New York, NY 10065, USA; Center for Virology and Vaccine Research, Beth Israel Deaconess Medical Center, Boston, MA 02215, USA; Howard Hughes Medical Institute

## Abstract

A clinical trial was performed to evaluate 3BNC117, a potent anti_HIV_1 antibody, in infected individuals during suppressive antiretroviral therapy (ART) and subsequent analytical treatment interruption (ATI). The circulating reservoir was evaluated by quantitative and qualitative outgrowth assay (Q^2^VOA) at entry and after 6 months, prior to ATI. Although there were no significant quantitative changes in the size of the reservoir, the composition of circulating reservoir clones varied over the 6_month period before treatment interruption in a manner that did not correlate with antibody sensitivity. The neutralization profile obtained from the reservoir by Q^2^VOA was predictive of time to rebound after ATI, and thus of antibody efficacy. Although 3BNC117 binding site amino acid variants found in rebound viruses pre_existed in the latent reservoir, only 3 of 217 rebound viruses were identical to 868 latent viruses. Instead many of the rebound viruses appeared to be recombinants, even in individuals with resistant reservoir viruses. By incorporating the possibility of recombination, 63% of the rebound viruses could have derived from the observed latent reservoir. In conclusion, viruses emerging during ATI in individuals treated with 3BNC117 are not the dominant species found in the circulating reservoir, but instead appear to represent recombinants.

**Summary:** In the setting of a clinical trial evaluating the anti_HIV_1 antibody 3BNC117, Cohen et al. demonstrate that rebound viruses that emerge following interruption of antiretroviral therapy are distinct from circulating latent viruses. However, rebound viruses often appear to be recombinants between isolated latent viruses.

## Introduction

Small molecule antiretroviral drugs are highly effective in suppressing HIV_1 viremia. However, therapy needs to be lifelong because it fails to eliminate a reservoir of latent HIV_1 viruses integrated into the genome of infected cells (Chun et al., 1997; Finzi et al., 1997). Significant efforts are currently focused on therapies, including immunotherapies, to target the reservoir to achieve sustainable ART_free remission (Churchill et al., 2016; Martin and Siliciano, 2016).

The immunotherapeutic agents that are clinically most advanced in this respect are newly discovered broad and potent monoclonal antibodies (bNAbs) that recognize the HIV_1 envelope protein expressed on the surface of infected cells and virions (Halper-Stromberg and Nussenzweig, 2016). These new antibodies protect against and suppress infection in mice and macaques (Barouch et al., 2013; Gautam et al., 2016; Halper-Stromberg et al., 2014; Horwitz et al., 2013; Klein et al., 2012; Shingai et al., 2014; Shingai et al., 2013). In human clinical trials, they suppress viremia, and delay viral rebound in the setting of treatment interruption (Bar et al., 2016; Caskey et al., 2015; Caskey et al., 2017; Lynch et al., 2015; Scheid et al., 2016). Most importantly, immunotherapy with bNAbs differs from small molecule drugs in that antibodies can eliminate circulating virus and infected cells through Fc_mediated effector mechanisms (Halper-Stromberg et al., 2014; Igarashi et al., 1999; Lu et al., 2016). In addition, bNAb administration is associated with development of potent antiviral CD8^+^ T cell immunity in macaques (Nishimura et al., 2017).

Infusion of VRC01, an anti_CD4_binding site antibody, in the setting of continued ART did not measurably alter the size of the latent reservoir in 6 individuals (Lynch et al., 2015). However,the sensitivity of circulating reservoir viruses to VRC01 was not determined and the relationship of latent viruses to plasma viruses that emerge during an analytical treatment interruption (ATI) was not examined.

Here we evaluate the effects of 3BNC117, a broad and potent anti_CD4_binding site bNAb (Caskey et al., 2015; Scheid et al., 2016; Scheid et al., 2011) in the setting of continued ART administration and during treatment interruption. We report on the dynamics of resistant and sensitive viruses in the latent HIV reservoir over a 6_month period prior to ATI, and the relationship between latent and rebound viruses.

## Results

### Study Participants

Fifteen HIV_1_infected participants virologically suppressed on ART were enrolled and underwent ATI (Table 1, Supplementary Table 1, and Supplementary Fig. 1). Participants received four intravenous infusions of 3BNC117 at 30 mg/kg at week 0, week 12, week 24, and week 27 (Fig. 1a). Leukapheresis was performed at week _2 and week 23 to collect PBMC for analyses of the latent reservoir. ART was discontinued 2 days after the week 24 infusion. Participants were not screened for 3BNC117 sensitivity before enrollment. All participants had a viral load of less than 50 copies/ml at day 0. The median baseline CD4^+^ T cell count was 688 cells/mm^3^, with a range of 391_1418 cells/mm^3^. Most participants entered the study on a non_ nucleoside reverse_transcriptase inhibitor (NNRTI) based ART regimen. Consistent with prior observations, 10 of the 15 participants (67%) had baseline bulk outgrowth culture viruses with 3BNC117 IC_50_ titers < 2.0 μg/ml (Supplementary Table 1) (Cohen et al., 2017; Scheid et al., 2016).

**Table 1.** Characteristics of participants at baseline (N=15)

|  |  |
| --- | --- |
| **Male sex** - n (%) | 14 (93) |
| **Median age** (range) | 43 (26-58) |
| **Race or ethnicity** |  |
| White non-Hispanic | 5 |
| Black non-Hispanic | 6 |
| Hispanic, regardless of race | 3 |
| Multiple, non-Hispanic | 1 |
| **HIV RNA <50 copies/ml (day 0)** – n (%) | 15 (100) |
| **CD4 T-cell count** – cells/mm<sup>3</sup> |  |
| Median (range) | 688 (391-1418) |
| **Nadir CD4 T-cell count** – cells/mm<sup>3</sup> |  |
| Median (range) | 350 (250-500) |
| **Years since starting ART** |  |
| Median (range) | 11 (1-21) |
| **Years on uninterrupted ART** |  |
| Median (range) | 7 (2-16) |
| **ART regimen** – n (%) |  |
| Integrase inhibitor based | 5 (33) |
| Protease inhibitor based | 2 (13) |
| NNRTI based | 8 (53) |
| **3BNC117 IC<sub>50</sub> < 2.0 µg/ml** – n (%) | 10 (67) |

**Figure 1.**
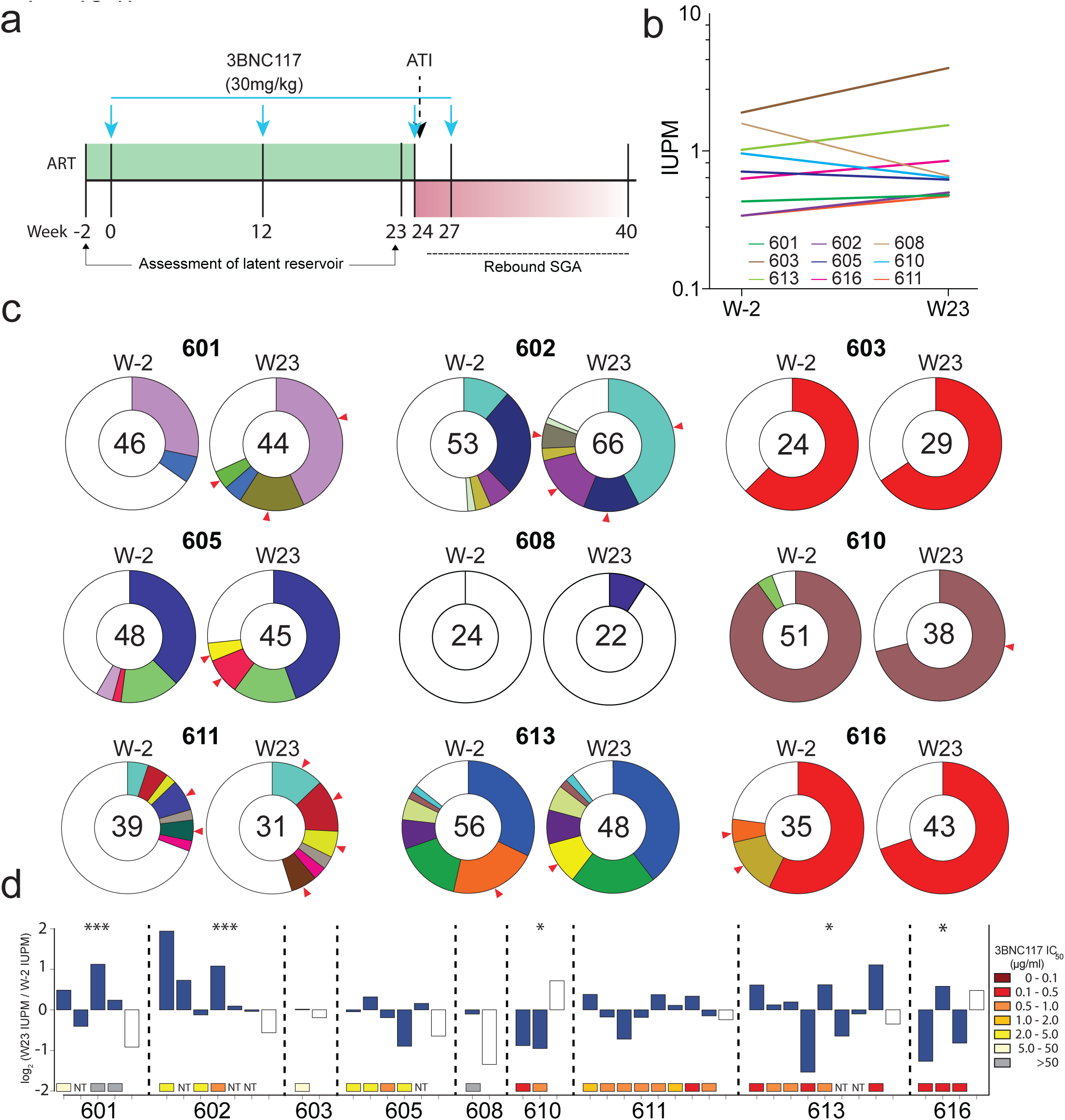
Changes in the circulating latent reservoir prior to treatment interruption. **a**, Study design, blue arrows represent 3BNC117 infusions. **b**, Change in infectious units per million (IUPM) of HIV_1 between week _2 and week 23, as determined by Q^2^VOA. There were no significant changes in IUPM between these two time points for any participant (defined by a > 6_fold change (Crooks et al., 2015)). **c**, Pie charts depicting the distribution of culture_derived *env* sequences from the 2 time points. The number in the inner circle indicates the total number of *env* sequences analyzed. White represents sequences isolated only once across both time points (singles), and colored areas represent identical sequences that appear more than once (clones). The size of the pie slice is proportional to the size of the clone. Red arrows denote clones that change in size between the 2 time points. **d,** Fluctuations within the latent reservoirs of each participant. Changes are measured in log_2_ fold change of IUPM between week _2 and week 23. Full bars represent clones and empty bars represent all singles as one group. *P* values from Fisher’s exact test (two_sided) are shown for participants with significant changes between the 2 time points. *** represents p_values < 0.001, * represents p_values < 0.05. Colored boxes below each bar represent the 3BNC117 IC_50_ titer of the particular clone. NT = Not tested.

### Safety and Pharmacokinetics

Infusions of 3BNC117 were generally well tolerated. One serious adverse event deemed not related to 3BNC117 occurred in a participant who required hospitalization for a psychotic episode associated with a new diagnosis of bipolar disorder. This participant was subsequently removed from the study before entering the ATI phase. Fourteen of the 15 participants who completed the study received all 4 infusions of 3BNC117. One participant received only 3 infusions because viral load results demonstrating viral rebound were available before the fourth infusion. Twenty_nine adverse events were considered at least possibly related to 3BNC117. Among these, 3 were graded as moderate and 1 was graded as severe. Complete data on adverse events is provided in Supplementary Table 2. In many participants, CD4^+^ T cell counts transiently declined during the ATI period (Supplementary Table 3, Supplementary Fig. 2). The adverse event profile observed here is similar to what has been reported in prior trials with 3BNC117 (Caskey et al., 2015; Scheid et al., 2016). The average half_life of 3BNC117 was 14.7 days, slightly shorter than the half_life of 17 days in HIV_uninfected individuals, but very similar to that previously reported for individuals receiving ART (Caskey et al., 2015; Scheid et al., 2016).

**Figure 2.**
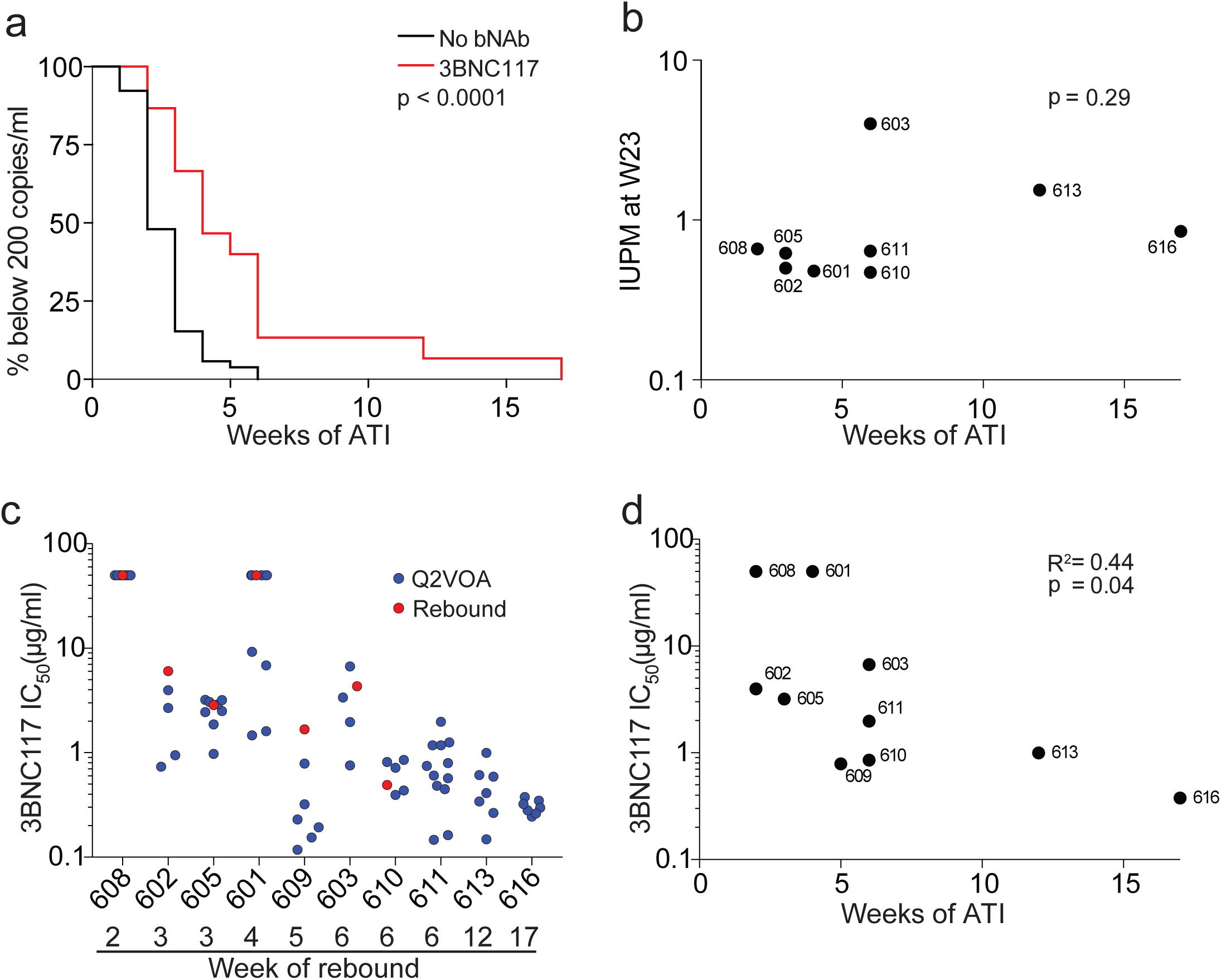
Time to rebound following treatment interruption. **a,** Kaplan_Meier plot summarizing viral rebound in 52 ACTG trial participants who underwent ATI without antibody treatment (black), and the 15 participants (red) who underwent ATI with 3BNC117 infusions. Y_ axis indicates percentage of participants with viral loads below 200 RNA copies/ml, x_axis indicates weeks after ATI. The p_value is based on the Log_rank test (two_sided). **b**, Dot plot representing the relationship between IUPM at week 23 and time to rebound for all participants for whom Q^2^VOA was performed. The p_value was calculated using the Spearman correlation (two_sided). **c**, Dot plot representing the 3BNC117 sensitivities of latent and rebound viruses.Q^2^VOA_derived latent viruses are shown as blue circles and culture_derived rebound viruses as red circles. Culture_derived rebound viruses could not be obtained from participants 611, 613, and 616. **d**, Dot plot representing the correlation between 3BNC117 sensitivity of the most resistant virus isolated by Q^2^VOA, and the week of rebound. The p_value was calculated using the Spearman correlation (two_sided).

### Size of the latent reservoir

The quantitative and qualitative viral outgrowth assay (Q^2^VOA) was used to examine the effect of 3BNC117 on the latent reservoir (Lorenzi et al., 2016). CD4^+^ T lymphocytes were collected by leukapheresis at weeks _2 and 23, during which time the participants remained on suppressive ART and received 2 infusions of 3BNC117 at weeks 0 and 12, maintaining detectable 3BNC117 serum levels for most of the 23_week period. Q^2^VOA was performed for 10 participants: the first 9 to enter the ATI phase of the trial (with the exception of participant 604, whose virus could not be grown in Q^2^VOA), and participant 616. Nine of these participants had samples available from both week _2 and week 23 (participant 609 declined the week 23 leukapheresis).

The infectious units per million (IUPM) of latently infected CD4^+^ T cells among these nine participants at weeks _2 and 23 ranged from 0.33_1.9 and 0.47_4.0, respectively. We found no significant changes in IUPM between the two time points for any participant (defined by a > 6_ fold change (Crooks et al., 2015) (Fig. 1b)). We conclude that 2 doses of 3BNC117 administered over 23 weeks in the setting of suppressive ART does not measurably reduce the size of the latent reservoir.

### The latent reservoir is dynamic over a 6_month period

To examine the molecular nature of the replication_competent viruses in the latent reservoir, we analyzed the latent viruses isolated by Q^2^VOA at weeks _2 and 23. An average of 1.18 ×; 10^8^ CD4^+^ T lymphocytes were cultured per participant per time point, yielding an average of 40 independent *env* sequences per participant per time point (Supplementary Table 4). Phylogenetic analysis showed that each participant analyzed was infected with epidemiologically unrelated clade B viruses (Supplementary Fig. 3). The clonal structure of the latent reservoir differed among the participants analyzed and could be divided into three categories: dominated by a single clone (603, 610, 616), dominated by multiple clones (601, 602, 605, 611, 613), or non_clonal (608) (Fig. 1c). Overall, 57% of viral sequences in the reservoir were clonal, which is consistent with prior studies (Bui et al., 2017; Hosmane et al., 2017; Lee et al., 2017; Lorenzi et al., 2016).

**Figure 3.**
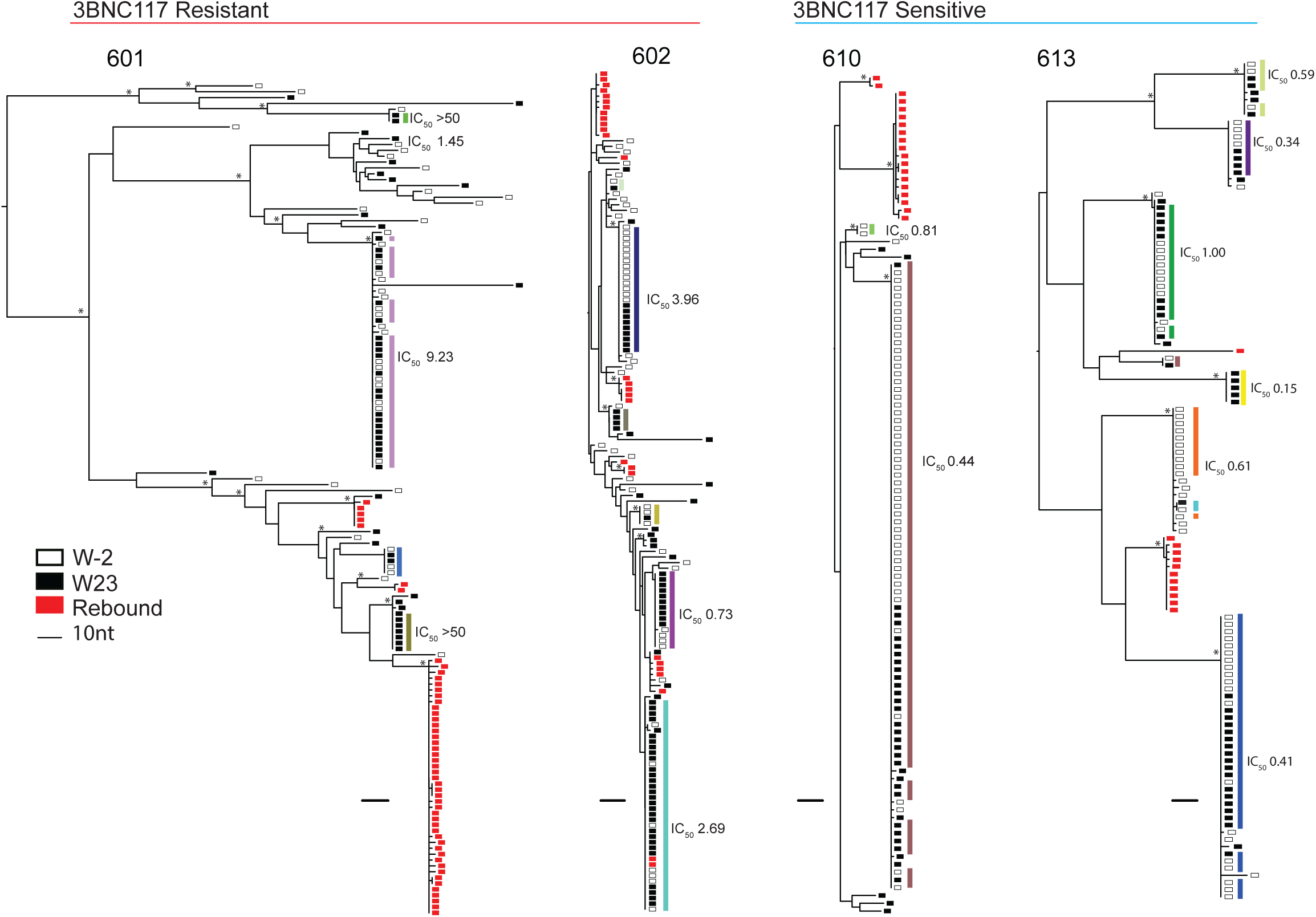
Phylogenetic trees of latent and rebound *env* sequences. Maximum likelihood phylogenetic trees of full_length *env* sequences of viruses isolated from Q^2^VOA outgrowth cultures and rebound SGA. Four participants with varying 3BNC117 sensitivity, clonal structure,and diversity are shown. Participants are categorized as 3BNC117 resistant or sensitive based on the presence of any resistant (IC_50_ > 2.0 μg/ml) viruses within the reservoir. Q^2^VOA_derived viruses from week _2 are represented as empty black rectangles, viruses from week 23 as full black rectangles, and rebound SGA viruses as red rectangles. Asterisks indicate nodes with significant bootstrap values (bootstrap support ≥90%). Clones are denoted by colored rectangles beside the phylogenetic tree. These colors correspond to colors in the pie charts in Figure 1c. Numbers represent 3BNC117 IC_50_ neutralization titers.

We observed clonal fluctuations that varied from appearance or disappearance of clones to large alterations in their relative size. For example, a clone from participant 602 comprised 11% of the circulating reservoir at week _2, but 42% of the circulating reservoir at week 23 (Fig. 1c). Utilizing the Q^2^VOA data and Bayesian inference, we inferred the IUPM of each clone at each time point. In 5 of the 9 participants, there were statistically significant fluctuations within the circulating reservoir, indicating that the observed changes in the composition of the reservoir between the 2 time points was very unlikely to be the result of finite sampling alone (Fig. 1d). However, these fluctuations were not correlated with the sensitivity of the outgrown viruses to 3BNC117, with some resistant clones decreasing in size, and some sensitive clones increasing in size (Fig. 1d). We conclude that the size of latent reservoir clones in circulation fluctuates over a 6_month interval in individuals treated with 3BNC117 while on suppressive ART, and that this effect is independent of bNAb sensitivity.

### 3BNC117 delays viral rebound in sensitive participants

Participants received 2 additional doses of 3BCN117 at weeks 24 and 27 in the setting of ART interruption (Fig. 1a). Viral rebound occurred between 2 and 17 weeks thereafter (Supplementary Fig. 4). The average time to rebound was 5.5 weeks, which was a significant delay compared to historical controls (p < 0.0001) (Fig 2a). The size of the latent reservoir, as determined by Q^2^VOA at week 23, did not correlate with time to rebound (Fig. 2b).

**Figure 4.**
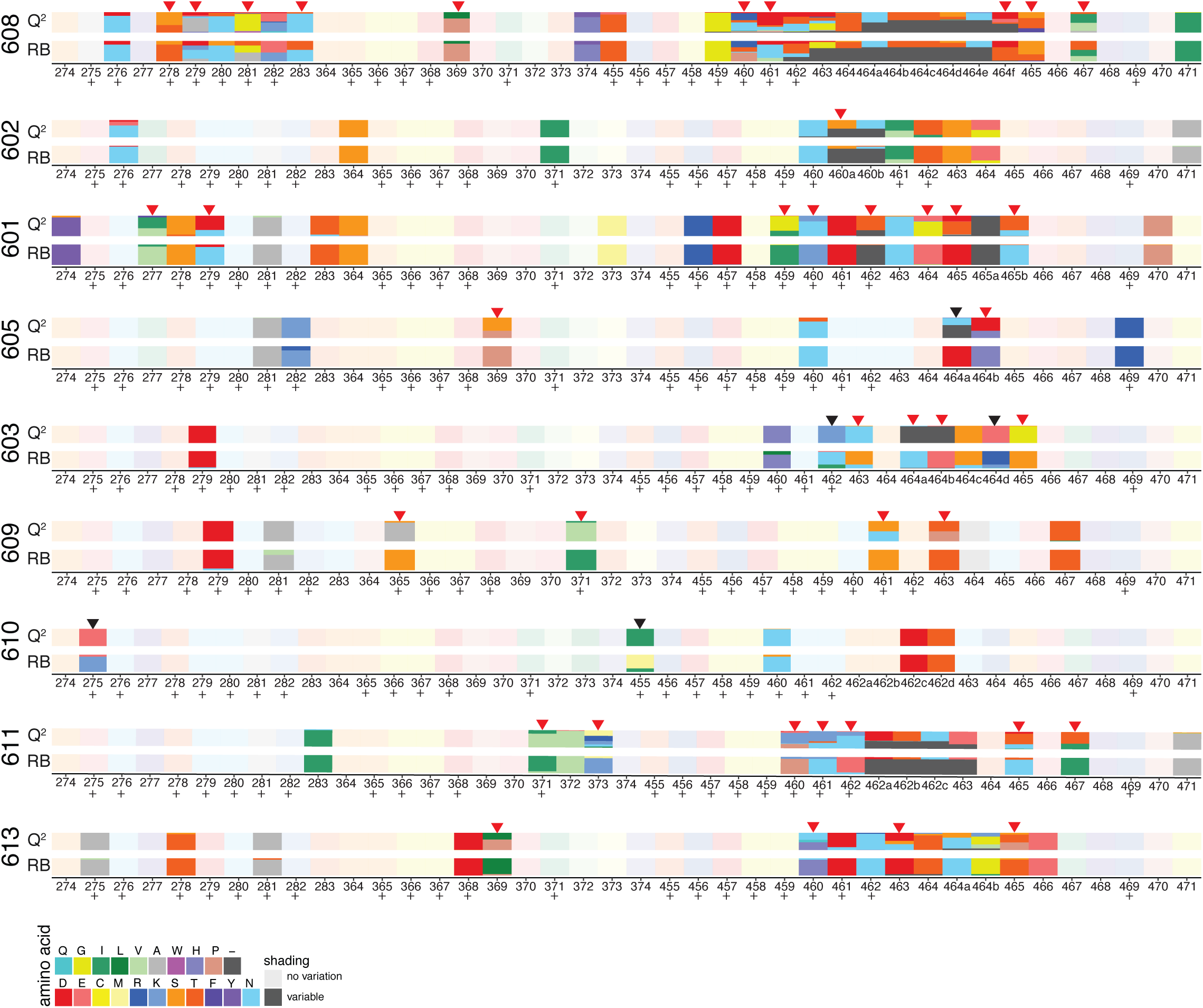
Amino acid variants at 3BNC117 binding sites in latent and rebound viruses. Chart illustrates amino acid changes in and around known 3BNC117 contact residues in Env (amino acid positions 274_283, 364_374, and 455 to 471), according to HXB2 numbering. + symbols represent 3BNC117 contact sites confirmed by crystal structures. Q^2^ indicates latent viruses isolated by Q^2^VOA, and RB represents rebound viruses isolated by SGA. Each amino acid is represented by a color, and the frequency of each amino acid is indicated by the height of the rectangle. Red arrows represent instances in which an amino acid found at low frequency in latent viruses became a majority amino acid in rebound viruses, and black arrows represent instances in which a majority rebound virus amino acid variant was not found in latent viruses.

For many participants, latent viruses exhibited a range of sensitives to 3BNC117 and other bNAbs that target distinct epitopes, as previously shown (Lorenzi et al., 2016) (Supplementary Tables 5 and 6). For example, viruses obtained by Q^2^VOA for participant 601 ranged from sensitive (IC_50_ = 1.45 μg/ml) to completely resistant (IC_50_ > 50 μg/ml) (Fig. 2c, blue circles). Participant 616, who demonstrated the longest delay in time to rebound, harbored a reservoir that was both highly sensitive and of restricted neutralization diversity (Fig. 2c). The 3BNC117 IC_50_ titer of the most resistant latent virus isolated from each of these participants correlated with time to rebound (p = 0.04, R^2^ = 0.44) (Fig. 2d). Thus, pre_existing resistance within the latent reservoir is predictive of bNAb efficacy.

To assess the neutralization sensitivity of rebound viruses, single genome analysis (SGA) was performed on plasma, and outgrowth cultures were performed using PBMCs obtained at the time of rebound. For 7 participants, outgrown viruses from the time of rebound were identical to plasma SGA viruses (Supplementary Fig. 5). Neutralization titers were then determined for the rebound viruses isolated from culture (Cohen et al., 2017). Overall, the 3BNC117 IC_50_ titers measured on the rebound viruses were similar to those found among the corresponding individual’s latent viruses (Fig. 2c, red circles). We conclude that the circulating latent viruses isolated by Q^2^VOA are representative of the overall neutralization profile present within the latent reservoir.

**Figure 5.**
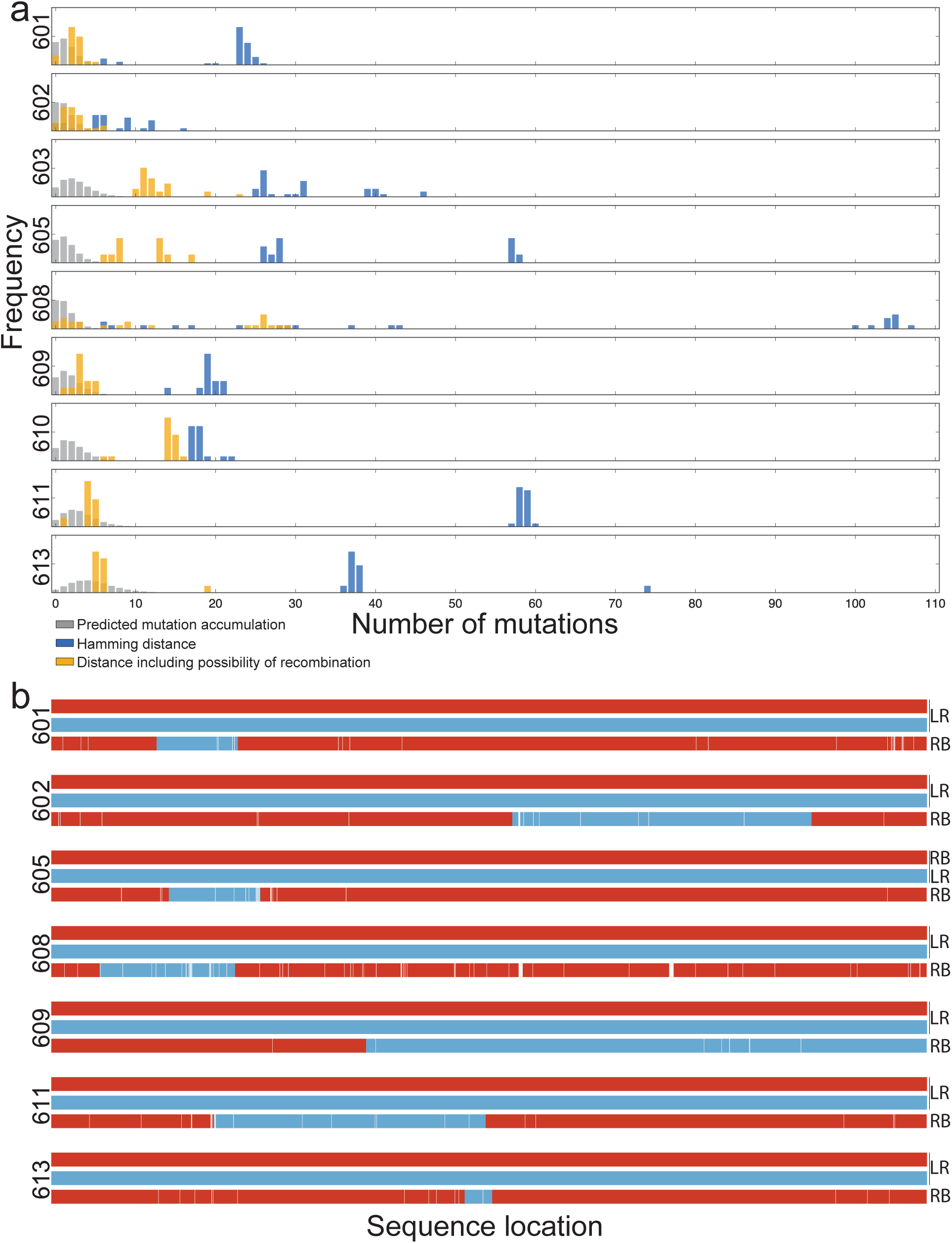
Relationship between latent and rebound sequences. **a,** Histograms show the percentage of *env* sequences (y_axis) and nucleotide distance (x_axis). The blue bars represent the observed Hamming distance between rebound and latent viruses. The grey bars represent the predicted distance between rebound and latent viruses based on a simulation of mutation accumulation during the ATI period for each participant. The yellow bars represent the minimal possible distance between the rebound and latent viruses including the possibility of recombination. **b,** Examples of putative recombination events. Parent *env* sequences are represented in red or blue and child sequences in combined red and blue. LR denotes latent reservoir sequences and RB denotes rebound sequences. The recombination breakpoints shown are the midpoints of the ranges determined by the 3SEQ algorithm (Supplementary Table 8). White lines represent point mutations unique to the child sequence. Parent and child sequences shown here are marked with green and red stars, respectively, in the phylogenetic trees in Supplementary Fig. 7.

### Rebound and latent viruses are distinct

Plasma rebound viruses isolated by SGA were compared to Q^2^VOA_derived latent viruses. Although some rebound viruses were closely related to reservoir viruses in resistant participants, only 3 of 217 rebound *env* sequences obtained were identical to the 768 Q^2^VOA_derived latent viruses isolated from 9 participants (Fig. 3 and Supplementary Fig 6). Moreover, the 3 exceptions were all found in a single individual (participant 602) and represented only 9.7% of all rebound sequences isolated from that participant (Fig. 3). Although the majority of analyzed participants harbored reservoirs dominated by expanded clones, these clones did not emerge in plasma upon treatment interruption. Strikingly, even a latent expanded clone with complete resistance to 3BNC117 could not be found among rebound viruses (Fig 3, participant 601).

### 3BNC117 resistance mutations in the latent reservoir

We compared the sequences of putative 3BNC117 target sites in *env* obtained from plasma rebound viruses to Q^2^VOA_derived sequences from the circulating reservoir for the presence of mutations (Bar et al., 2016; Caskey et al., 2015; Cheng et al., 2018; Lynch et al., 2015; Scheid et al., 2016). Despite the lack of overlap between the rebound and latent viruses noted above, rebound viruses demonstrated nucleotide and corresponding amino acid variants in the 3BNC117 binding site that preexisted in the latent reservoir. These variants were often present at low frequencies in the reservoir and enriched in rebound viruses (Figure 4, red arrows). For example, all rebound sequences from participant 601 contain an isoleucine at position 459, compared to 33 percent of sequences within the latent reservoir. In participant 609, all rebound sequences at position 371 contain a leucine, which is present in just a small minority of latent reservoir sequences. On the other hand, there were only 5 cases in which amino acid variants found in the majority of rebound sequences were not detected in the latent reservoir (Figure 4, black arrows). We conclude that while latent and rebound viruses are distinct, 3BNC117 binding site variants present in rebound viruses are frequently pre_existing within the observed latent reservoir.

### Relationship between rebound viruses and circulating latent viruses

Despite the fact that 3BNC117 binding site variant amino acids found in rebound sequences were already present within the latent reservoir, identical reservoir and rebound sequences were exceedingly rare. While occasional rebound viruses were very similar to latent viruses, the average Hamming distance between rebound and latent sequences among all participants was 31 nucleotides, excluding insertions and deletions.

Since Q^2^VOA is limited by its dependence on reactivation of latent viruses *in vitro* (Bruner et al., 2016; Ho et al., 2013; Hosmane et al., 2017) we also examined latent viruses by a reactivation_ independent method. Near full_length HIV genome analysis (Bruner et al., 2016; Ho et al., 2013) was performed on week 23 samples for participants with Q^2^VOA viruses that displayed varying sensitivity, phylogenetic diversity, and clonal structure. Between 9 – 33 near full_length sequences were obtained from each of 6 participants, for a total of 133 sequences. For participants 603 and 616, whose reservoirs are dominated by a large clone, 85% of near full_ length sequence *env* genes were identical to Q^2^VOA_derived viruses. For the other 4 participants, the identity between near_full length sequences and Q^2^VOA_derived viruses ranged from 8.6% to 47.3% (Supplementary Table 4). However, we found no instance of identical near full_length HIV sequences and rebound viruses among the 5 participants for whom rebound SGA sequences were available (Supplementary Fig. 7). The average Hamming distance between near full_length HIV sequences and rebound viruses was 32 nucleotides, excluding insertions and deletions, which is similar to that for Q^2^VOA_derived sequences. We conclude that intact proviral DNA sequences obtained from circulating CD4^+^ T cells overlap with sequences obtained by Q^2^VOA but differ from sequences obtained from rebound plasma SGA.

To examine the accumulation of mutations during ATI as a potential explanation for the divergence between latent and rebound viruses, we created a mathematical model to simulate this process (Supplementary Fig. 8). For each participant we computed the frequency of mutations accumulated during the simulated rebound, where the time to rebound in simulation was within one week before or after the observed time to rebound for that participant (Fig. 5a gray bars). Using this model and including 868 *env* sequences from both near full_length sequencing and Q^2^VOA (Fig. 5a blue bars), we found only 7 rebound sequences that were within the number of expected mutations, and all of these were found in participant 602. We then analyzed latent and rebound sequences for the presence of recombination using the 3SEQ recombination algorithm (http://mol.ax/software/3seq/). We found statistical evidence of recombination (rejection of the null hypothesis of clonal evolution) among 7 of the 9 participants tested (Supplementary Table 7). In the majority of recombination events, the 2 “parent”sequences derived from the latent reservoir, and the “child”sequence was a rebound sequence; however, there were instances in which the parents were comprised of either a latent and rebound sequence or two rebound sequences (Supplementary Fig. 7, and Supplementary Table 8). Examples of recombinant rebound viruses are illustrated in Figure 5b. Of note, there was only a single instance among the 27 latent parents involved in recombination events in which the latent virus was an expanded clone. In this case (participant 613), the rebound virus that derived from this recombination event accounted for only 1 out of the 12 rebound viruses isolated. The other 11 rebound viruses derived from a recombination event between two latent viruses that were “singles”(Supplementary Fig. 7 and Supplementary Tables 8 and 9). Taking into account the possibility of recombination, the observed distance between latent and rebound viruses decreases, and is often close to the range predicted by the mutation simulation model described above (Fig. 5a yellow bars). We conclude that in addition to the accumulation of mutations during ATI, recombination may account for the considerable genetic distance between circulating latent and rebound virus.

## Discussion

We enrolled 15 HIV_1_infected individuals to receive 2 doses of monoclonal antibody 3BNC117 over a 23_week period while on suppressive ART and to subsequently undergo ART interruption while receiving 2 additional doses of the antibody. Our analysis revealed that the size of the circulating reservoir is not altered during the 23_week observation period before ART interruption. However, the composition of the circulating latent reservoir varies significantly, in a way that is independent of the sensitivity of the latent viruses to 3BNC117. The neutralization profile obtained from the reservoir by Q^2^VOA is predictive of time to rebound, and rebound virus amino acid variants are frequently pre_existing within the latent reservoir. Finally, although the rebounding viruses are typically absent from the circulating latent reservoir, the sequences of rebound viruses frequently correspond to latent virus recombinants.

Antibody monotherapy in the absence of ART delays viral rebound and selects for antibody_ resistant viruses (Caskey et al., 2015; Caskey et al., 2017; Scheid et al., 2016). When participants in a prior trial were pre_screened for sensitivity and received infusions of 3BNC117 at time 0 and 3 weeks, rebound was delayed for an average of 6.7 weeks (Scheid et al., 2016). The same regimen was administered in the setting of the ATI in the current study, but in participants who were not pre_screened for sensitivity. Rebound was delayed for an average of 5.5 weeks, and an analysis of the reservoir by Q^2^VOA revealed that the delay in rebound by 3BNC117 monotherapy is directly related to the sensitivity of T cell derived outgrowth viruses found in the circulating reservoir. Thus, the outgrown viruses obtained by Q^2^VOA are representative of the neutralization profile of the clinically relevant latent reservoir.

The size of the latent reservoir remains relatively constant over time, with a calculated half_life of 3.6 years (Crooks et al., 2015; Finzi et al., 1999). However, the discovery of replication_ competent clones suggested a dynamic reservoir (Cohn and Nussenzweig, 2017; Kwon and Siliciano, 2017), and clones of T cells bearing integrated viruses have been reported to expand and contract over periods of years (Cohn et al., 2015; Wang et al., 2018). Expression of viral *env* in individuals on suppressive ART is limited, and neither 3BNC117 nor VRC01 (Lynch et al., 2015) infusion altered the size of the HIV reservoir in this setting. Nevertheless, there was significant fluctuation in the distribution of circulating latent viruses over the 6_month period before ART interruption that was not directly related to 3BNC117 sensitivity. These changes could be due to homeostatic proliferation, stimulation by cognate antigen, or cell death. For example, an influenza_specific CD4^+^ T cell clone that harbors a latent virus might expand after vaccination or exposure to the pathogen and later contract. Alternatively, the observed differences could simply be due to changes in the specific T cell clones that happen to be in circulation at the 2 time points. Our finding that the distribution of circulating clones of latent cells can change dramatically in a relatively short period of 6 months indicates that the circulating reservoir is far more dynamic than previously appreciated.

Viral outgrowth cultures utilizing peripheral blood mononuclear cells are limited in that they only capture the circulating fraction of the reservoir that can be reactivated *in vitro*. Nevertheless, even with limited sampling in 6 participants, there was a 48.7% overall overlap between 133 near full_length provirus sequences amplified from DNA, and 513 sequences obtained from viruses emerging in Q^2^VOA (Supplementary Table 4, Supplementary Fig. 6). Thus, there is good concordance between the near full_length DNA assay and Q^2^VOA.

Although we obtained an average of 80 latent HIV_1 sequences for each participant, and 57% of these were clonal, we only found three instances in which the rebounding virus was identical to a virus in the latent reservoir. However, 3BNC117 binding site nucleotide and amino acid variants found in rebound sequences were frequently preexisting in the latent reservoir. The observations that rebound viruses are unique but contain specific mutations found in the reservoir appear to be at odds but can be explained at least in part by recombination between viruses.

Recombination is a major source of HIV_1 diversity (Burke, 1997; Robertson et al., 1995). Recombination occurs when two RNA strands from different viruses co_infect a single cell, wherein recombinant genomes are generated by reverse transcriptase template switching. HIV_1 recombination rates have been estimated at ˜1.4 ×; 10^−5^ to 1.38 ×; 10^−4^ events per base per generation (Batorsky et al., 2011; Neher and Leitner, 2010; Shriner et al., 2004). This rate is within range of that of reported HIV mutation rates (Abram et al., 2010; Ji and Loeb, 1992; Roberts et al., 1988). Recombination has been found to mediate escape from ART (Kellam and Larder, 1995; Moutouh et al., 1996), CD8^+^ T cells (Ritchie et al., 2014; Streeck et al., 2008), and autologous neutralizing antibodies (Chaillon et al., 2013; Moore et al., 2013; Song et al., 2018).

By incorporating the possibility of recombination and estimating the number of mutations that might have occurred during the ATI period for each participant, approximately 63% (137 of 217) of rebounding viruses could have derived from the observed latent reservoir. This includes 5 out of 9 individuals in which all or nearly all of the rebound viruses could be accounted for by this approach. Thus, in 56% of the individuals studied, the latent viruses detected by Q^2^VOA or near full_length genome sequencing appeared to have contributed genetic information to the viruses that emerge at the time of rebound. Recombinants were found in 3BNC117_sensitive participants and also in the participant with complete resistance to 3BNC117 (608), and therefore these events do not appear to be dependent on bNAb administration. While expanded clones comprised 57% of all outgrowth sequences, in only a single instance did a parent recombinant sequence come from an expanded latent clone. This could be because expanded T cell clones harbor viruses with reduced fitness, or because of differences in reactivation requirements of latent viruses *in vivo* and *in vitro*, such that latent viruses in large clones readily reactivate under *in vitro* conditions but are more resistant to reactivation *in vivo*. The latter would be consistent with the ability of the clones to expand without undergoing apoptosis or pyroptosis, as typically happens during productive infection. Finally, the number of recombination events observed likely represents an underestimate since we only considered recombination events within *env*.

It has been suggested that recombination may be important for the survival of reactivated latent viruses, which could exhibit decreased fitness either as a distinguishing characteristic of their ability to remain in a latent state (de Verneuil et al., 2018) or due to increased sensitivity to the host immune response (Immonen et al., 2015). While our findings suggest that recombination does in fact confer a fitness advantage, our data do not allow us to determine when recombination occurred. If the recombinants are preexisting but not detected in circulation before the rebound, it would imply that they are either extremely rare in circulation, or that circulating T cells are not representative of the reservoir. If recombination occurs at the time of rebound, it may have significant implications for understanding the frequency with which latent viruses are reactivated.

In conclusion, by examining the effects of antibody therapy on the circulating reservoir we document its dynamic nature and its relationship to viruses that emerge upon interruption of antiretroviral therapy in individuals receiving 3BNC117.

## Materials and Methods

### Study Design

An open label study was conducted in HIV_infected participants virologically suppressed on ART (clinicaltrials.gov NCT02588586). Study participants were enrolled sequentially according to eligibility criteria. All participants provided written informed consent and the study was conducted in accordance with International Conference on Harmonisation Good Clinical Practice guidelines. The study protocol was approved by the Institutional Review Boards at The Rockefeller University and Weill Cornell Medical Center. Participants were followed for a total of 60 weeks from the time of enrollment. The primary outcome measures of the study were safety and the time to virologic rebound after ART interruption.

### Study Participants

Study participants were recruited at The Rockefeller University Hospital and Weill Cornell Medical Center Clinical Trials Unit, both in New York, USA. All infusions were performed at The Rockefeller University Hospital. Eligible participants were HIV_infected adults on ART aged 18_65 years with plasma HIV_1 RNA 50 copies/ml for 12 months prior to enrollment and < 20 copies/ml at the time of screening, and CD4 count > 500/μl. Exclusion criteria included hepatitis B or C infection, history of AIDS_defining illness within 1 year prior to enrollment, CD4 nadir < 200/μl, and significant medical conditions or laboratory abnormalities. Participants on an NNRTI were switched to an integrase inhibitor_based regimen 4 weeks before treatment interruption to avoid monotherapy as a result of the prolonged half_life of NNRTIs. Time to viral rebound was compared to a historical cohort of 52 participants who underwent ATI without intervention in trials performed by the ACTG, as previously described(Scheid et al., 2016).

### Study Procedures

The appropriate volume of 3BNC117 was calculated according to body weight, diluted in sterile normal saline to a total volume of 250 ml, and administered intravenously over 60 min. Study participants received 30mg/kg of 3BNC117 on weeks 0, 12, 24, and 27 and remained under monitoring at The Rockefeller University Hospital for 4 hours after each infusion. Participants returned for frequent follow up visits for safety assessments, which included physical examination, measurement of clinical laboratory parameters such as hematology, chemistries, urinalysis, and pregnancy tests (for women). Leukapheresis was performed at The Rockefeller University Hospital at week _2 and week 23. ART treatment was interrupted 2 days after the third 3BNC117 infusion (week 24). ART was reinitiated after two consecutive plasma viral load measurements exceeded 200 copies/ml or CD4 counts fell below 350/μl. Plasma HIV_1 RNA levels were monitored weekly during the ATI period and CD4 counts were measured every other week. Study investigators evaluated and graded adverse events according to the DAIDS AE Grading Table v2.0 and determined causality. The CTCAE v4.03 grading scale was used for reporting and grading adverse events related to infusion reactions. Blood samples were collected before and at multiple times after 3BNC117 infusion. Samples were processed within 4 hours of collection, and serum and plasma samples were stored at −80 °C. PBMCs were isolated by density gradient centrifugation. The absolute number of peripheral blood mononuclear cells was determined by an automated cell counter (Vi_Cell XR; Beckman Coulter), and cells were cryopreserved in fetal bovine serum plus 10% DMSO.

### Quantitative and qualitative viral outgrowth assay (Q^2^VOA)

The quantitative and qualitative viral outgrowth assay (Q^2^VOA) was performed utilizing PBMC isolated by leukapheresis, as previously described (Lorenzi et al., 2016). Q^2^VOA isolates replication_competent viruses from the latent reservoir using a limiting dilution method such that each virus likely originates from a single reactivated infectious provirus. The frequency of latently infected cells was calculated using the IUPMStats v1.0 (http://silicianolab.johnshopkins.edu) (Rosenbloom et al., 2015).

### Viral neutralization testing by TZM.bl neutralization assay

Supernatants from p24 positive Q^2^VOA wells were tested against a panel of broadly neutralizing antibodies by the TZM.bl neutralization assay, as described (Li et al., 2005; Montefiori, 2005). Neutralization assays were conducted in a laboratory meeting Good Clinical Laboratory Practice (GCLP) Quality Assurance criteria (Michael S. Seaman, Beth Israel Deaconess Medical Center).

### Measurement of 3BNC117 serum levels by TZM.bl neutralization assay

Serum concentrations of 3BNC117 were measured at multiple time points post_infusion using a validated luciferase_based neutralization assay in TZM.bl cells as previously described (Sarzotti-Kelsoe et al., 2014). Briefly, serum samples were tested using a primary 1:20 dilution with 5_ fold titration series against HIV_1 Env pseudovirus Q769.d22, which is highly sensitive to neutralization by 3BNC117. Env pseudoviruses were produced using an ART_resistant backbone vector that reduces background inhibitory activity of antiretroviral drugs if present in the serum sample (SG3ΔEnv/K101P.Q148H.Y181C, M. Seaman unpublished data). 3BNC117 clinical drug product was also tested in every assay set_up using a primary concentration of 10 μg/ml with 5_fold titration series. The serum concentration of 3BNC117 for each sample was calculated as follows: serum ID_50_ titer (dilution) x 3BNC117 IC50 titer (μg/ml) = serum concentration of 3BNC117 (μg/ml). Murine leukemia virus (MuLV) was utilized as a negative control. All assays were performed in a laboratory meeting GCLP standards. The half_life of 3BNC117 was calculated using Phoenix WinNonLin Build 8 (Certara).

### Bulk Cultures

We performed bulk viral outgrowth cultures using PBMCs harvested at week _2, and from rebound samples as described (Caskey et al., 2015). Sequence analysis on bulk culture viruses was performed as previously described (Lorenzi et al., 2016).

### Single_genome amplification (SGA) of plasma rebound virus *env* genes

Single_genome amplification and sequencing of HIV_1 *env* genes was performed as previously described (Caskey et al., 2017; Salazar-Gonzalez et al., 2008; Scheid et al., 2016).

### Q^2^VOA_derived virus sequence analysis

HIV *env* sequences from p24 positive supernatants were obtained and analyzed as previously described (Lorenzi et al., 2016). Phylogenetic analysis was performed by generating nucleotide alignments using MAFFT (Katoh and Standley, 2013) and posterior phylogenetic trees using PhyML(Guindon et al., 2010), both installed as plugins on Geneious software version 10.2 (Biomatters).

### Near full_length genome analysis

Genomic DNA was extracted from 1_10×10^6^ CD4^+^ T cells from week 23 leukapheresis samples. DNA was subjected to limiting_dilution PCR using semi_nested primers in the *gag* gene 3GagININ 5’_GGGGCTGTTGGCTCTGGT_3’(Bruner et al., 2016; Ho et al., 2013). PCR products were visualized and quantified using 1% 96_well E_Gels (Invitrogen). DNA dilutions wherein < 30% of the *gag* PCR wells were positive, were selected for further analysis because they have a > 90% of probability of containing single copy of HIV DNA in each PCR reaction based on the Poisson distribution. Near full_length outer PCR was performed and 1μl aliquots subjected to nested *env* PCR (Ho et al., 2013; Li et al., 2007). Samples containing approximately 2600bp *env* amplicons were subjected to four segment PCR (A, B, C, D) (Ho et al., 2013). Samples containing either segment A+C, or A+D, or B+C, or B+D were subjected to library preparation and sequencing (Lorenzi et al., 2016). Sequence adapters were removed using Cutadapt v1.9.1 and read assembly for each virus was performed in three steps: 1. de novo assembly was performed using Spades v3.9.0 to yield long contig files; 2. Contigs longer than 255bp were aligned to an HIV full genome reference sequence and a consensus sequence was generated using Mira assembly v4.0.2; 3. Reads were re_aligned to the consensus sequence to close gaps, and a final read consensus was generated for each sequence. Sequences with double peaks (cutoff consensus identity for any residue <75%), stop codons, or shorter than the expected near full genome size were omitted from downstream analyses.

### 3BNC117 binding site analysis

All *env* sequences were translated to amino acids and aligned using ClustalW (Larkin et al., 2007). The HXB2 (K03455) *env* sequence was used as a reference for numbering the amino acids. Frequency plots were produced for each *env* position targeted by 3BNC117 (Zhou et al., 2013).

### Clonal fluctuation calculation

Clonal composition shifts between time points in individual participants. We employed a statistical test to gauge whether such changes in composition could be explained by finite sampling alone, or whether they indicated real changes in the relative proportions of each clone in the reservoir. For this analysis, the counts for all sequences that were only observed once across time points were merged together in a group of “singles.”All other sequence counts were considered separately. We used Fisher’s exact test, implemented in R, to determine whether the counts of each clone (including the singles) observed at each time point were consistent with a single underlying distribution for both time points. IUPM were inferred for each clone based on the particular Q^2^VOA run from which each member of the clone was isolated, as previously described (Lorenzi et al., 2016). Bayesian Markov chain Monte Carlo was implemented in Stan, using four separate chains of 10^5^ iterations. The first 50000 iterations for each chain were discarded as warm_up.

### Simulation of mutation accumulation during rebound

Rebounding viral sequences were rarely identical to those observed in viral outgrowth assays or by near full_length sequencing. To understand whether the differences between rebound sequences and those in the observed reservoir might be attributed to mutations accumulated during the rebound process, or whether the rebound virus was instead likely to have originated from clones that were not previously observed we developed a stochastic mutation simulation model.

The model is based on that of Hill and collaborators (Hill et al., 2014). We begin with a population of 10^6^ latently infected cells. Latent viruses can reactivate with rate *A*, in which case the latently infected cell becomes actively infected. The product of these two parameters (the number of latently infected cells and the reactivation rate) gives the approximate number of reactivation events per day. Actively infected cells die with rate *d*, without infecting other cells, or burst with rate *b*. In the latter case the actively infected cell also dies, but the release of virions results in the active infection of *c* new cells. During each burst event, the number of new infected cells *c* is assumed to follow a Poisson distribution. In principle, latently infected cells could also die or homeostatically expand, but we assume that such processes typically occur at much slower rates than the fast replication dynamics of actively_infected cells. Therefore, they are not expected to contribute substantially to the rebound process, so we do not include these possibilities in our simulation.

The underlying parameters (reactivation rate, death rate of actively infected cells, etc.) are not perfectly constrained by experiment, and may vary between individuals. Following the approach of Hill *et al*. (Hill et al., 2014), we therefore choose the net growth rate of rebound virus, the ratio of the variance to the mean number of new infected cells during a cell bursting event, and the rate of reactivation of latently infected cells, to be random within the bounds of experimental constraints (see Hill *et al*. for further details). In addition to these parameters, we also allow the net death rate of actively infected cells to vary, which was not included in the analysis of Hill *et al*. We use the results of Markowitz *et al*. (Markowitz et al., 2003) to constrain the death rate of infected cells, which was estimated to be approximately 1 day^−1^ with a standard deviation of 0.3 day^−1^.

In addition to this simple model tracking the number of latent and actively infected cells, we also consider evolution of the virus at the sequence level. In order to compare with the data obtained in this study, we assume a sequence length for *Env* of 2600 base pairs. We are specifically interested in the accumulation of *de novo* mutations during the rebound process. We therefore treat the length 2600 sequences as binary, where zeros represent nucleotides that are the same as the source sequence from the latent reservoir, and ones represent a mutation. This approximation is justified because the number of mutations observed is much smaller than the length of the sequence, and thus the probability of back mutations is very small. We assume that mutations occur during the reverse transcription process, when new infection events occur after an actively infected cell bursts. The probability of mutation per site per new infection event is 3×10^−5^ (Sanjuan et al., 2010). In addition we allow for the possibility of recombination between sequences, with a probability of crossover between strands of 1.4×10^−5^ per site per new infection event (Neher and Leitner, 2010). This probability also implicitly includes the probability of coinfection of a single host cell.

For simplicity, we assume that mutations have no effect on the fitness (or replication rate) of the virus. However, it is known that nonsynonymous mutations are often strongly deleterious, including in HIV (Haddox et al., 2016; Loeb et al., 1989; Louie et al., 2018; Zanini et al., 2017). Thus, our assumption of neutrality likely overestimates the true number of mutations that would be accumulated during the rebound process.

We simulate this stochastic rebound process until either the number of actively infected cells reaches 3×10^5^, corresponding roughly to a threshold of detection for virus in the blood around 200 copies mL^−1^ (Hill et al., 2014), or until the total simulation time reaches 100 days. We repeated these simulations using 10^4^ random sets of parameters as described above. Different parameter choices and stochastic effects lead to a range of observed rebound times (Supplementary Fig. 8). At the end of each simulation, we also record the distribution of the number of mutations accumulated during the rebound process. As expected, the average number of accumulated mutations in the rebound sequences increases with the rebound time, though there is substantial stochastic variation (Supplementary Fig. 8).

### Distance, recombination snippet

In order to conservatively estimate the true distance (in terms of number of mutations) between rebound sequences and those in the reservoir, we counted each observed variant in rebound sequences as a mutation only if it did not match with any of the nucleotides at that site in the observed reservoir sequences. This prevents us from computing spuriously large mutational distances simply due to recombination between diverse sequences in the reservoir during the rebound process. The estimated rate of recombination in HIV is high, comparable to the mutation rate (Neher and Leitner, 2010), and thus the possibility of recombination cannot be simply neglected. Indeed, tests for signatures of recombination (Lam et al., 2018) in the rebound sequences found multiple likely recombination events for all patients in this study except for 603 and 610.

## Code availability

Code used for these simulations is available on GitHub at https://github.com/bartonlab/simple_ rebound_simulation.

## Data availability

Sequence data that support the findings of this study have been deposited in GenBank with the accession codes pending.

## Acknowledgements

We thank all study participants who devoted time to our research, and members of the Nussenzweig lab, especially Lilian Cohn, Yotam Bar_On, Till Schoofs, and Jill Horowitz for helpful discussions. We also thank Steve Smiley, Randy Tressler, Pat Fast, and Harriet Park.

## Funding

Y.Z.C. is supported by grant KL2 TR001865 from the National Center for Advancing Translational Sciences (NCATS) and grant UL1 TR000043 from the National Institutes of Health (NIH) Clinical and Translational Science Award (CTSA) program. This work was supported by NIH/National Institute of Allergy and Infectious Diseases grant U01 AI118536 (M.C.). This work was supported in part by Bill and Melinda Gates Foundation Collaboration for AIDS Vaccine Discovery (CAVD) grants OPP1092074 and OPP1124068, the NIH Center for HIV/AIDS Vaccine Immunology and Immunogen Discovery (CHAVI_ID) (1UM1 AI100663) (M.C.N.); a BEAT_HIV Delaney grant UM1 AI126620 (M.C.), the Einstein_Rockefeller_CUNY Center for AIDS Research (1P30AI124414_01A1) and the Robertson fund. M.C.N. is a Howard Hughes Medical Institute Investigator.

## Author Contributions

Y.Z.C., J.C.C.L., L.B., R.M.G., M.C., and M.C.N. designed the research. Y.Z.C., J.C.C.L, L.K., L.B., C_L.L., P.M.D., K.M., A.B., J.P.D., S.A.B., M.W_P., I.S., M.S.S., and M.C. performed the research. Y.Z.C., J.C.C.L., L.K., J.P.B., J.P., M.C., and M.C.N. analyzed the data. Y.Z.C., J.C.C.L., J.P.B., M.C., and M.C.N. wrote the manuscript.

## Competing Interests

There is a patent on 3BNC117 on which M.C.N is an inventor.

## Abbreviations

ART: Antiretroviral therapy
ATI: Analytical treatment interruption
bNAb: Broadly neutralizing antibody
IUPM: Infectious units per million
Q^2^VOA: Quantitative and qualitative viral outgrowth assay
NNRTI: Non_nucleoside reverse_transcriptase inhibitor
SGA: Single genome analysis

